# β-arrestin-mediated Angiotensin II type 1 Receptor Activation Promotes Pulmonary Vascular Remodeling in Pulmonary Hypertension

**DOI:** 10.1101/2021.03.23.436676

**Authors:** Zhiyuan Ma, Gayathri Viswanathan, Mason Sellig, Chanpreet Jassal, Issac Choi, Xinyu Xiong, Nour Nazo, Sudarshan Rajagopal

**Affiliations:** Division of Cardiology, Department of Medicine, Duke University School of Medicine, Durham, NC, USA; Trinity College of Arts and Sciences, Duke University, Durham, NC, USA; The University of North Carolina, Chapel Hill, NC, USA; Department of Biochemistry, Duke University Medical Center, Durham, NC, USA

**Keywords:** Pulmonary arterial hypertension, Angiotensin, G protein-coupled receptor, Biased agonism, Beta-arrestin

## Abstract

**Objectives:** The goal of this study was to test whether a β-arrestin-biased agonist of the angiotensin II (AngII) type 1 receptor (AT_1_R), which acts as a vasodilator while not blocking cellular proliferation, would have positive effects compared to a balanced agonist, angiotensin II (AngII), or an antagonist, losartan, in pulmonary arterial hypertension (PAH).

**Background:** PAH is a disease of abnormal pulmonary vascular remodeling whose treatment has focused on targeting vasoactive substances, such as inhibiting endothelin signaling and promoting prostacyclin signaling. PAH medical therapies are thought to primarily act as vasodilators, although they may also have effects on pulmonary vascular remodeling. There are a number of reports that blocking AT_1_R signaling can be beneficial in preclinical models of PAH. The AT_1_R is a G protein-coupled receptor (GPCR) that promotes vasoconstriction through heterotrimeric G proteins but also signals via β-arrestins, which promote cardioprotective effects and vasodilation.

**Methods:** We compared the effects of a β-arrestin-biased AT1R agonist, TRV120023 (TRV023), to a balanced agonist (AngII) and an antagonist (losartan) in preclinical PAH models.

**Results:** In acute infusion studies, AngII increased right ventricular (RV) pressures while TRV023 did not. However, with chronic infusion in monocrotaline (MCT) PAH rats, TRV023 failed to improve hemodynamics or survival compared to AngII, while losartan significantly improved survival. Both TRV023 and AngII enhanced proliferation and migration of pulmonary artery smooth muscle cells (PASMCs) from PAH patients, which was associated with the promotion of proliferative MAP kinase signaling.

**Conclusions:** β-arrestin-mediated AT_1_R signaling promotes vascular remodeling and worsens PAH, and suggests that the primary benefit of current PAH therapies is through pulmonary vascular reverse remodeling and not vasodilation.

## Introduction

Pulmonary arterial hypertension (PAH) is a disease associated with excessive pulmonary vascular remodeling characterized by dysfunction of endothelial cells, proliferation of smooth muscle cells that leads obliteration of pulmonary arterioles. This results in a high pulmonary vascular resistance, right ventricular (RV) hypertrophy, dilation and ultimately failure (1). Current PAH therapies largely target vasoactive mediators such as endothelin-1 and prostacyclin, which signal through G protein-coupled receptors such as the type A endothelin receptor (ET_A_R) and prostacyclin receptor (IP), respectively (2). However, the utility of these therapies is thought to be limited in their efficacy by acting primarily as pulmonary vasodilators and not affecting the pulmonary vascular remodeling that underlies progression of the disease (3). Other vasoactive mediators, such as angiotensin II (AngII), are also thought to contribute to the pathobiology of PAH. The expression of AngII and AT_1_R is elevated in PAH and increased levels of AngII are associated with PAH progression and mortality (4–7). Blocking AT_1_R signaling is beneficial in rats with PAH, including restored RV dysfunction, decreased pulmonary vascular remodeling, and delayed PAH progression (4,8).

GPCRs such as the AT_1_R signal canonically through heterotrimeric G proteins and multifunctional β-arrestin adapter proteins, which in addition to signaling, also promote receptor internalization and desensitization of G protein signaling (9). Subsequently, G protein-dependent (10) and/or β-arrestin-dependent (11) signaling is transduced by the activation of downstream mediators, such as Ca^2+^ -dependent PKC, MAPK signaling, and GPCR-mediated transactivation of receptor tyrosine kinases (12,13). G protein-mediated AT_1_R signaling has been found to lead to hypertension, myocardial hypertrophy, and cardiac dysfunction (13,14), while β-arrestin-mediated signaling has been shown to promote cardioprotective effects and decrease systemic and renovascular resistance in systolic heart failure (15–18). Recently, “biased” AT_1_R ligands to engage and stabilize distinct active conformations of AT_1_R (19) and activate only a subset of receptor-mediated signaling pathways (15,20), such as only through G proteins (“G protein-biased”) or β-arrestins (“β-arrestin-biased”) (21,22). For example, compounds such as TRV120027 and TRV120023 (TRV023) have been shown to function as β-arrestin-biased agonists, antagonizing G protein signaling and activate β-arrestin-mediated signaling. These drugs act acutely act as vasodilators, by antagonizing AT_1_R G protein-mediated vasoconstriction, while promoting cardiac function and decreasing apoptosis through their β-arrest-mediated signaling.

It is not known whether the benefit of current therapies is primarily mediated through their action vasodilation or reverse remodeling of the pulmonary vasculature, as patients on long-standing PAH therapies can still display severe vascular pathology (23). We hypothesized that we could address this question through the selective activation of AT_1_R β-arrestin-mediated signaling with a β-arrestin-biased agonist, which would be predicted to act as an acute vasodilator while promoting AT_1_R β-arrestin-mediated signaling. We compared the physiological and molecular effects of a β-arrestin-biased AT_1_R agonist, TRV120023 (TRV023), to a balanced agonist, AngII, and an antagonist, losartan, in the treatment of PAH. We found that the β-arrestin-biased AT_1_R stimulation promoted pulmonary vascular remodeling, demonstrating that vasodilation in the presence of β-arrestin-biased AT_1_R activation worsens PAH. These findings suggest that the beneficial long-term effects of drugs targeting GPCR signaling in PAH is not through vasodilation but from reversing pulmonary vascular remodeling.

## Methods

### AT_1_R ligands

AngII and TRV023 (Sar-Arg-Val-Tyr-Lys-His-Pro-Ala-OH) were synthesized by GenScript USA (Piscataway, NJ) with quality control assessed by high-performance chromatography and mass spectrometry. Losartan was purchased from Santa Cruz biotechnology (sc-204796). AngII was also obtained from Sigma (A9525-1MG).

### Animals

All animal experiments were conducted in compliance with institutional guidelines and were approved by Duke University Institutional Animal Care and Use Committee (Protocol: A175-16-08). Male Sprague–Dawley (SD) rats (5-6 weeks old, Charles River Laboratories) weighing between 150 and 200 grams were used in the chronic infusion studies. For acute infusion, 8 to 10-week old Sprague-Dawley rats with an average weight of 300 grams were used.

### Experimental protocols for PAH treatment

Pulmonary hypertension was induced by a single subcutaneous injection of crotaline (MCT, 60 mg·kg^−1^; Sigma). For chronic treatments, either vehicle (PBS), 1 mg·kg^−1^·day^−1^ AngII, 10 mg·kg^−1^ ·day^−1^ losartan, or 14.4 mg·kg^−1^ ·day^−1^ (10 µg·kg^−1^·min^−1^) TRV023 were delivered by Alzet osmotic mini-pumps (Model 2ML4 or 2ML2; Cupertino, CA) placed in a subcutaneous pouch under anesthesia with isoflurane. Osmotic pumps were implanted 2 weeks after MCT injection for a duration of 4 weeks for survival studies and 1 week after MCT injection for a duration of 2 weeks for histology and hemodynamic analyses.

### Hemodynamic Analysis

For acute hemodynamic responses, the level of anesthesia was regulated by delivery of 3% isoflurane administered through a nose cone with 100% O_2_. Rats were placed on a servo‐controlled heating table to maintain rectal temperature constant at 37°C. A midline skin incision was made to expose the trachea, carotid artery and jugular vein. A pressure transducer-tipped curved 2 French catheter (SPR-513, Millar) was inserted into the right ventricle via right internal jugular vein access, a pressure-volume catheter was inserted into the aorta and left-ventricle via the left carotid artery (SPR-869, Millar), and the left internal jugular vein was accessed for the administration of intravenous fluids and drugs. After obtaining vascular access, the animal was allowed to stabilize for 15 min before the beginning of the experimental protocol. Rats (n=6) were infused with increasing doses of AngII or TRV023 followed by treatment with losartan. Dosages of AngII and TRV023 were determined based on previous studies (24,25). At 5-minute intervals, either increasing doses (1, 10, and 100 µg/kg) of AngII or increasing doses (10, 100 µg/kg, 1mg/kg) of TRV023 were administered followed by treatment with 100 µg/kg losartan via the left internal jugular vein at a volume of 1 µl/g body wt (25–30 µl total volume) followed immediately by 30 µl of normal saline. Before the injection of vasoactive agents, each rat received an equivalent volume (55–60 µl, 2 μl/g body wt) of normal saline as a vehicle control. Peak rates of LV and RV pressure rise (dP/dtmax) and fall (dP/dtmin) were determined. RV and LV pressures were measured at end-diastole (EDRVP and EDLVP, respectively) and at peak-systole (RVPmax and LVPmax). After the procedure, the rats were euthanized by bilateral thoracotomy. Data was analyzed in LabChart (ADI Instruments).

For chronic hemodynamic responses, the open-chest approach for RV heart catheterization was performed as previously described (26). Rats were anesthetized with xylazine (2.5 mg/kg) and ketamine (10 mg/kg). A needle was inserted into the trachea to serve as endotracheal intubation, the cannula connected to a volume cycled rodent ventilator on normal air with a tidal volume of 1.2 ml and respiratory rate of 82/min. The conductance catheter tip was inserted through a stab wound on the apical RV free wall until all electrodes were inside the ventricle. RV preloads was altered by tightening a suture placed around the IVC to perform RV pressure-volume (PV) loop recordings.

Right ventricular hypertrophy was quantified as the ratio of right ventricular to body weight (RV/BW). Similarly, left ventricular hypertrophy was determined as the ratio of left ventricular and septal weight to body weight ((LV+S)/BW). Measurements were performed by investigators blinded to the experimental groups.

### Immunofluorescence and Morphometric Analysis

Immunofluorescence and morphometric quantification were perform as previously reported (27). Briefly, lung tissues were paraffin-embedded and sectioned at 5 μm. Sections were incubated at 4°C with anti-VWF (1:200, Dako) and anti-α-smooth muscle actin (1:100, Sigma) antibodies in 10% goat norm serum in PBS overnight, followed by incubation with AlexaFluor 488 goat anti-mouse or AlexaFluor 594 goat anti-rabbit secondary antibody for 1 h in the dark. DNA was stained with Hoechst. Images were acquired using a LSM upright 780 confocal microscope. For the morphometric analyses, ten random fields were examined for 20-80 Lm muscular arteries. The external and internal media perimeters of muscular arteries were measured using ImageJ and external and internal media radii were calculated using r = perimeter/2π. The medial wall thickness was expressed as (external media radii – internal media radii)/external media radii. Quantifications were performed by investigators blinded to the experimental groups.

### Immunohistochemistry

Immunohistochemistry for lung tissues from MCT rats was performed by Department of Pathology, Duke University Medical Center. Briefly, antigen was retrieved at 100°Cfor 20 minutes in pH 6.1 citrate buffer. Lung sections were stained for Ki67 (RM-9106-S, 1:400, Thermo Scientific) and Cleaved Caspase-3 (9661s, 1:400, Cell Signaling Technology). To assess proliferative and apoptosis markers in pulmonary arteries of MCT rats treated with AngII or TRV023 or losartan, five random fields were examined for 50- to 80-μm arteries from 5 rats for each experimental group. Using ZEN lite software Ki67 and cleaved caspase-3 positive cells to total number of cell ratios were determined in the pulmonary arteries. Quantifications were performed by investigators blinded to the experimental groups.

### Human samples

Human pulmonary arterial smooth muscle cells (PASMCs) were isolated by culture from explanted lung tissue from confirmed subjects with PAH undergoing lung transplantation at Duke University Medical Center. The study protocol for human tissue donation was approved by the Institutional Review Board of Duke University Medical Center (Pro00044369) and written informed consent was obtained from each subject. Control pulmonary arterial smooth muscle cells were obtained from human lung tissues from normal donor lungs at the Cleveland Clinic (gift of Drs. Suzy Comhair and Serpil Erzurum) and from pooled primary cells from a commercial sources (Lonza).

Pulmonary arteries were minced and digested overnight in 30 ml of digesting solution (HBSS, collagenase (0.1 mg/ml), DNase (0.1 mg/ml), HEPES (2.5 ml), penicillin/streptomycin (250 mg/ml), amphotericin B (0.625 mg/ml) and 1%anti-anti). Cells were then filtered using 100μm-pore nylon cell strainer and neutralized with 10 ml SMC growth media. Cells were centrifuged at 330g for 7 mins and resuspended with SMC growth medium. Cells were cultured in SMC medium (310-500, Cell Applications) containing growth supplements (311-GS, Cell Applications), 1% anti-anti and 1% penicillin/streptomycin.

### BrdU proliferation assay

Cell proliferation was assessed by Cell Proliferation ELISA BrdU kit (11647229001, Roche) according to the manufacturer’s instructions. Around 8000 PASMCs were seeded in each well of 96-well plate. At 60-70% confluence, cells were starved overnight and stimulation with 100 nM AngII for 24 hours and 1 μM TRV023 for 48 hours. BrdU was added to the cells during last 6-8 hours of incubation. Cells were then fixed for 30 mins and probed with anti-BrdU-POD antibody for 90 minutes followed by substrate development. The chemiluminescence was quantified by measuring the absorbance at 370nm and 492nm using a BioTek Synergy Neo 2 multi-mode reader.

### In vitro Scratch Assay

PASMCs were seeded on 24-well plates. At 80-90% confluency cells were serum starved for 24 hours and scratches were made using a 20 μl pipette tip. Cells were washed with Dulbecco’s phosphate-buffered saline (PBS without calcium and magnesium) and stimulated with basal smooth muscle cell medium containing 1 μM AngII, 10 μM TRV023 with or without 10 μM losartan. The wound closure was monitored using a live-cell station Zeiss Axio Observer microscope (Duke Light Microscopy Core Facility). The images were captured in real-time at 0 hour and for every hour for 12 hours. The initial edges of the scratch at 0th hour was marked and migrated distance at 12 hours was measured using MetaMorph Premier (Molecular Devices).

### Protein Isolation and Immunoblotting

Human PASMCs were serum starved for 24 hours and treated with increasing concentration of AngII or TRV023. The lysates were collected after 10 and 20 mins after treatment using 1 X SDS buffer. The proteins were denatured at 100^°^C for 10 mins. The lysates were stored at −20°. Total proteins extracted with 1 X SDS were separated by 10% sodium dodecyl sulfate-polyacrylamide gel electrophoresis (SDS-PAGE) and transferred to nitrocellulose membranes (1620112, Bio-Rad). The membranes were blocked in 5% BSA for 1 hour at RT and probed with primary antibodies, ERK (06-182, Millipore Sigma), phospho-ERK (8201S, Cell Signaling), p38 (8690S, Cell Signaling), phospho-p38 (4511S, Cell Signaling) at 4°C overnight. The membranes were incubated with secondary antibodies for 1 hour at RT. The blots were developed using SuperSignal™ West Pico PLUS Chemiluminescent Substrate (34580, Thermo Fisher Scientific) and captured by Bio-Rad ChemiDoc Imaging System. The data analysis was performed using ImageLab software (Bio-Rad).

### Next-Generation Sequencing (RNA-seq)

Total RNA was extracted from ≈10 mg of MCT rat lung tissues using the RNeasy Fibrous Tissue Mini Kit (74704, Qiagen) following to the manufacturer’s instructions. Total RNA was submitted to Novogene Corporation, Inc., for quality analysis, RNA sequencing and bioinformatics analysis. All RNA samples had RNA integrity numbers ~ 8-9. All qualified samples were processed for library construction followed by sequencing. Reads were aligned to the rat genome and differential gene expression analysis (DEG) were obtained. The Database for Annotation, Visualization and Integrated Discovery (DAVID) v6.8 was used to analyze the signaling pathways using *Rattus norvegicus* as background.

### RNA isolation

The PASMCs isolated from PH patients were treated with 500nM AngII and 5μM TRV023 for 4 hours. The total RNA from PASMCs were isolated by using Qiagen RNeasy Plus Mini/Micro kit. The procedures were followed according to the manufacturer’s instructions. Finally, RNA was dissolved in DEPC-water and stored at −80°C. The quality and concentration of total RNA was measured by NanoDrop spectrophotometer. Total RNA of 1μg was reverse transcribed into cDNA using Bio-Rad iScript cDNA synthesis kit according to the manufacturer’s instructions.

### Quantitative polymerase chain reaction (qPCR)

The cDNA was mixed with iTaqTM SYBR® Green Supermix and the relevant primers. The PCR primers for 18S rRNA, MMP-1, MMP-2, MMP-7, MMP-9, TIMP-1, TIMP-2, TIMP-3 and TIMP-4 are shown in Table I supplement. The cDNA levels were measured using the Applied Biosystems 7300 Real-Time PCR System. The PCR was performed by initial incubation at 50°C for 2 min, denaturing at 95°C for 30 seconds, then 40 cycles of denaturing at 95°C for 15 sec and annealing at 60°C for 1 min. Data are expressed as the relative expression (fold change = 2−∆∆Ct) of each target gene compared to 18S rRNA.

### Statistics

All graphs and data generated in this study were analyzed using GraphPad Prism 8 Software. All quantitative data is presented as means ± SEM. The statistical significance of differences was determined using Student’s two-tailed t test in two groups, and one-way or two-way ANOVA along with *Bonferroni, Tukey’s* or *Sidak’s* multiple comparison test in multiple groups. Survival curves were analyzed by the Kaplan-Meier method and compared by a log-rank test. A p value < 0.05 was considered statistically significant.

## Results

### Acute infusion of β-arrestin-biased AT_1_R agonist does not increase LV or RV pressure

Upon stimulation by the endogenous ligand AngII, AT_1_R activates both heterotrimeric G proteins and β-arrestin adapter proteins. Stimulation with the β-arrestin-biased agonist TRV023 has been reported to only robustly recruit β-arrestin in the absence of G protein activation, enhancing myocyte contractility but without promoting hypertrophy as seen with AngII (24). Since AngII and TRV023 activate distinct signaling pathways through AT_1_R, we first set out to examine the physiological effects of AngII or TRV023 on LV and RV hemodynamics with acute infusion of drug in healthy Sprague-Dawley rats while monitoring their hemodynamics with simultaneous right and left heart catheterization. Acute infusion of AngII in normal rats increased both LV pressure and contractility (dP/dt) while TRV023 did not (**Fig. 1A** and **B**). Similarly, AngII increased RV pressure while TRV023 did not (**Fig. 1C**), while neither drug had a significant effect on RV contractility (**Fig. 1D**). Notably, neither AngII or TRV023 had a significant effect on LV filling pressure (data not shown). These findings are consistent with G protein-mediated AT_1_R signaling promoting both systemic and pulmonary vasoconstriction. Importantly, blocking both β-arrestin and G-protein-mediated AT_1_R signaling by the administration of losartan completely reversed AngII-induced elevated LV pressure.

**Figure 1.**
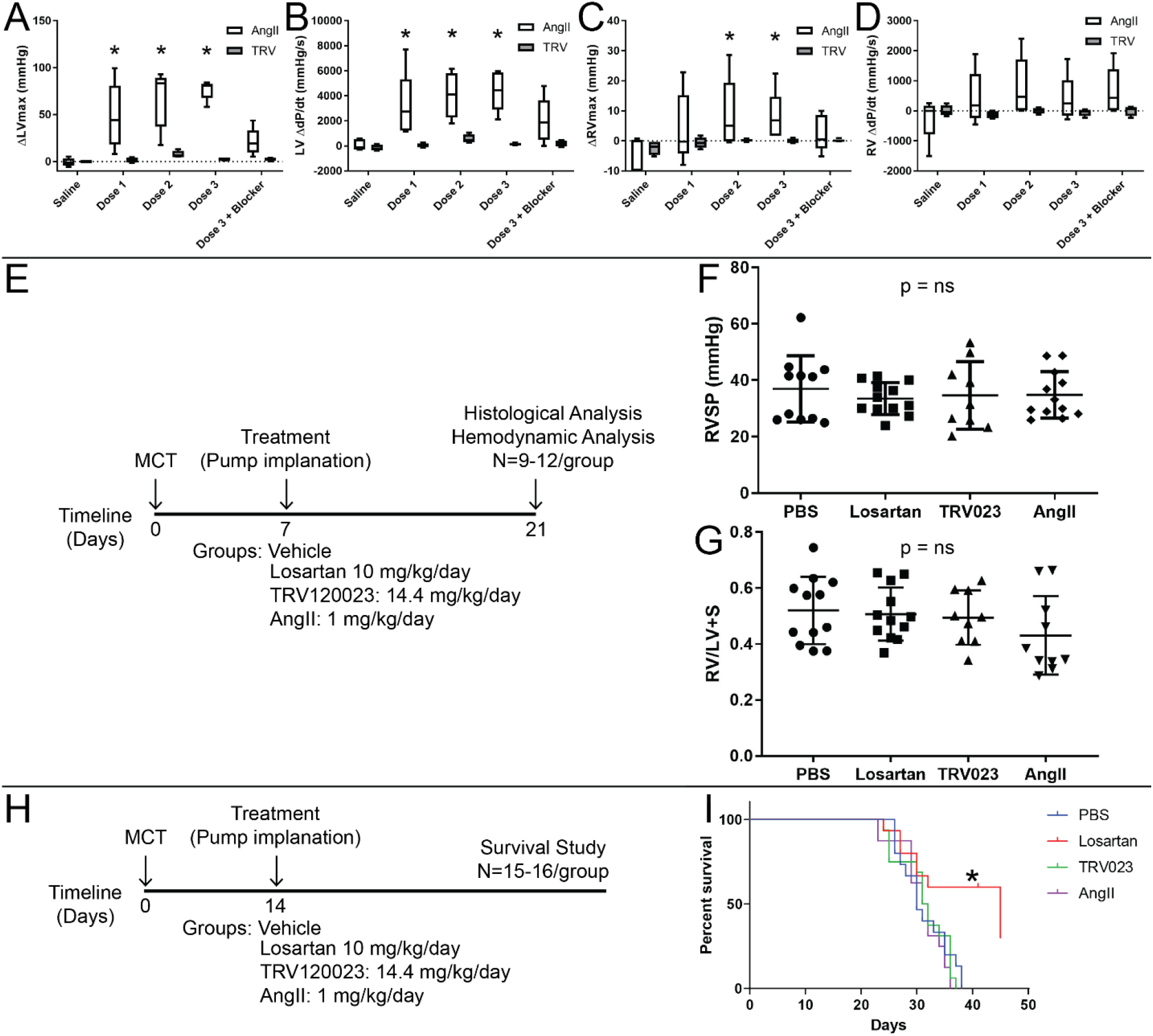
Hemodynamic effects of targeting AT_1_R with acute or chronic infusions in healthy and MCT PH rats. Acute infusion of AngII in healthy Sprague-Dawley rats resulted in significant increases in left ventricular (LV) (**A**) pressure and (**B**) contractility (dP/dt – change in pressure over change in time) while TRV023 did not. AngII infusion also increased right ventricular (RV) (**C**) pressure but not (**D**) contractility, while TRV023 did not have a significant effect on either. With chronic infusion in MCT PH rats, one week after MCT injection a (**E**) 2 week infusion of AngII, TRV023, losartan or vehicle did not result in significant changes in (**F**) right ventricular systolic pressure (RVSP) or (**G**) RV hypertrophy as assessed by Fulton index (right ventricle (RV) mass / left ventricle (LV) + septum (S) mass). Assessing survival by Kaplan-Meier in MCT PH rats (**H**), two weeks after MCT injection (**I**) an infusion of losartan had a significant effect on survival, while the AngII, TRV023 and vehicle infusions had no effect.

### β-arrestin-mediated AT_1_R activation does not improve hemodynamics in PAH rats in chronic infusion

To determine how β-arrestin-biased AT_1_R signaling affects hemodynamics in PAH, we treated MCT PAH rats 1 week after MCT injection with PBS, AngII, TRV023 or losartan for 2 weeks followed by an analysis of hemodynamics with RV PV loop analysis (**Fig. 1E**). In contrast to acute infusion, chronic treatment of MCT PAH rats with TRV023 or losartan did not result in any significant improvement in RV systolic pressure (RVSP) compared to either PBS or AngII treatment (**Fig. 1F**). Similarly, there were no significant differences in most hemodynamic parameters from PV loop analysis (**Supp. Fig. 1**). Since there was no effect of chronic infusion of AngII, TRV023 or losartan on RV hemodynamics in MCT PAH rats, we next assessed RV and LV hypertrophy. We found that LV hypertrophy was increased in rats that received TRV023 or AngII compared with losartan (**Supp. Fig. 2**), consistent with the significant effect of AngII on LV hemodynamics, while there were no differences in RV hypertrophy among treatment groups (**Fig. 1G**). While this was largely consistent with previous studies demonstrating no significant hemodynamic benefit of AT_1_R antagonism in MCT PAH (4), it suggested that the previously described beneficial effects of AT_1_R antagonism in PAH were not directly mediated through an improvement in RV hemodynamics.

### Blocking both β-arrestin and G protein-mediated AT1R signaling improves survival in MCT PAH

To determine whether β-arrestin-mediated AT_1_R activation had beneficial effects on survival, rats were injected with MCT to induce PAH, and then treated with AngII, TRV023 or losartan for 4 weeks via minipump (**Fig. 1H**). Compared to vehicle-treated group, losartan treatment markedly improved survival compared to AngII, TRV023 and PBS (**Fig. 1I**). Notably, AngII treatment did not worsen survival relative to PBS, which is likely due to high levels of AT_1_R activation in vehicle conditions. These results demonstrate that antagonizing both β-arrestin and G protein-dependent AT_1_R signaling is beneficial in PAH, but the selective inhibition of G protein-dependent signaling with a β-arrestin-biased agonist (TRV023) did not improve hemodynamics or outcomes in PAH. This suggests that β-arrestin-mediated AT_1_R signaling contributes to PAH pathology.

### β-arrestin-mediated AT_1_R signaling stimulates cell proliferation in vivo

As we did not observe significant hemodynamic changes between groups although there was a significant difference in survival, we next examined the pathology of lungs in MCT PAH rats treated with AngII, TRV023, losartan and PBS. Consistent with no changes in the measured RV hemodynamics, there were no significant differences in the medial wall thickness of small pulmonary arteries in rats treated with PBS, AngII, TRV023 or losartan (**Fig. 2A** and **B**). However, Ki67-positive cells in vascular walls were significantly increased in rats that received TRV023 compared to PBS or losartan (**Fig. 2C** and **D**). Moreover, AngII infusion significantly increased proapoptotic activity as assessed by cleaved caspase-3 staining compared to control, but not compared to TRV023, which may partially explain why AngII infusion did not increase the number of Ki67 positive cells (**Fig. 2E** and **F**). Taken together, these findings suggest that β-arrestin-mediated AT_1_R signaling by TRV023 promotes specific aspects of adverse pulmonary vascular remodeling through stimulating cell proliferation, while the effects of AngII predominantly affect apoptosis.

**Figure 2.**
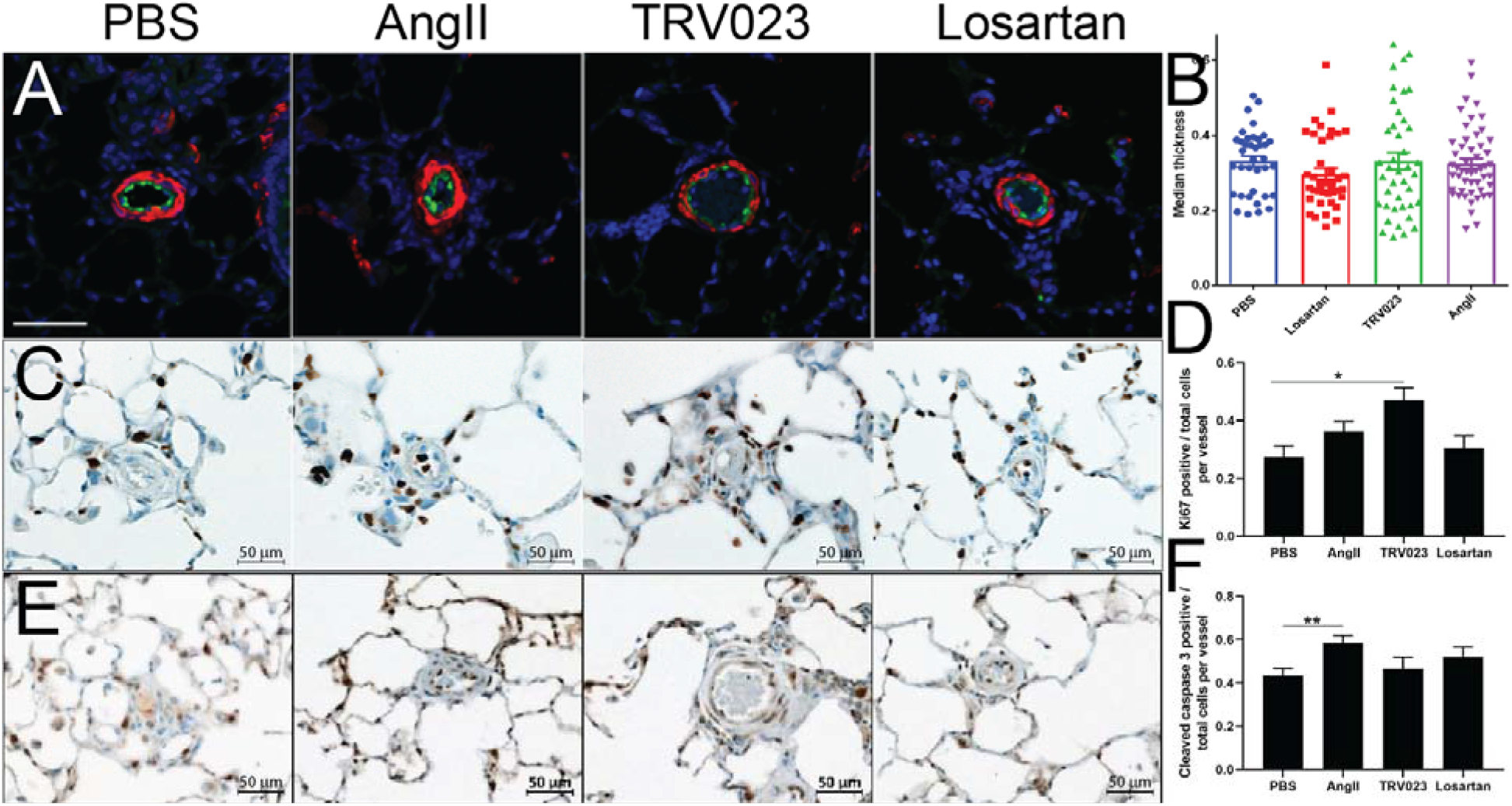
Effect of TRV023 on cellular proliferation in the lungs of MCT rats. Lungs from MCT rats treated with AngII, TRV023, losartan or vehicle were inflation-fixed in 10% formalin, paraffin-embedded and then sectioned for staining. (**A**) Endothelial cell staining with vWF (green) and α-smooth muscle cell actin (red) demonstrated (**B**) no significant differences in gross pulmonary vascular remodeling. For immunohistochemical staining with Ki67 and cleaved caspase-3, the number of Ki67/cleaved caspase-3 -positive (brown) and Ki67/cleaved caspase-3 -negative cells (blue) were counted in pulmonary arteries (PA) and the ratio of positive/total cells were calculated. Representative micrographs showing that (**C**) Ki67 and (**E**) cleaved caspase-3 -positive cells were stained in brown and the negative cells in blue. Quantitative assessment of (**D**) Ki67- and (**F**) cleaved caspase-3 -positive cells from MCT rats. n = 5 rats for each group. n=5 PA for each rat. Each bar represents Mean ± SEM (n = 25). **p < 0.005, **p < 0.05 versus vehicle treated MCT rats. ^##^p < 0,005 versus losartan treated MCT rats. Scale = 50 μm.

### AngII and TRV023 Activate Transcriptional Programs for Pulmonary Vascular Remodeling

To identify the signaling pathways that may underlie disease progression in AT_1_R ligand-treated rats, we performed lung transcriptome analysis from MCT rats treated with AngII, TRV023, losartan and PBS with RNA-seq. Comparing drug-treated to vehicle-treated groups, only 3 genes were differentially regulated between vehicle and losartan, compared to 93 genes differentially regulated between vehicle and TRV023, and 195 genes differentially regulated between vehicle and AngII (**Fig. 3A**). We performed gene ontology (GO) enrichment to identify specific cellular processes associated with these differentially regulated genes, which were consistent with TRV023 and AngII promoting changes in cell proliferation and cell energetics respectively (**Fig. 3B**). Only 13 differentially expressed transcripts were shared between the AngII and TRV023 groups (**Fig. 3C**), although hierarchical clustering of the differentially expressed genes demonstrated that the AngII and TRV023 groups were more similar to each other than to the losartan and vehicle groups (**Fig. 3D**). To identify the underlying pathways common to the 13 genes that were regulated in common in the AngII and TRV023 groups, we used the Database for Annotation, Visualization and Integrated Discovery (DAVID) bioinformatics database and PANTHER analysis. Among those 13 genes that were regulated in common between AngII and TRV023 (**Fig. 3E**), these cell signaling pathway enrichment analysis tools revealed upregulation of transcripts in AngII and TRV023 MCT lungs that are known to be regulated by mitogen-activated protein kinase (MAPK) signaling cascade, including triggering receptor expressed on myeloid cells 2 (Trem2) (28) and growth differentiation factor 15 (Gdf15) (29,30) (**Fig. 3F**). Activation/phosphorylation of MAPK isotypes, such as extracellular signal-regulated kinases (ERKs), mitogen activated protein kinase 14 (p-38), and mitogen-activated protein kinase 8 (JNK), are known to regulate in vascular smooth muscle cell proliferation, extracellular matrix production and migration (31–33). In our transcriptome analysis we also found a phosphatase inhibitor, cAMP-regulated phosphoprotein 19 (Arpp19) that is downregulated in both Angiotensin-2 and TRV023 MCT rat lungs (**Fig. 3F**).

**Figure 3.**
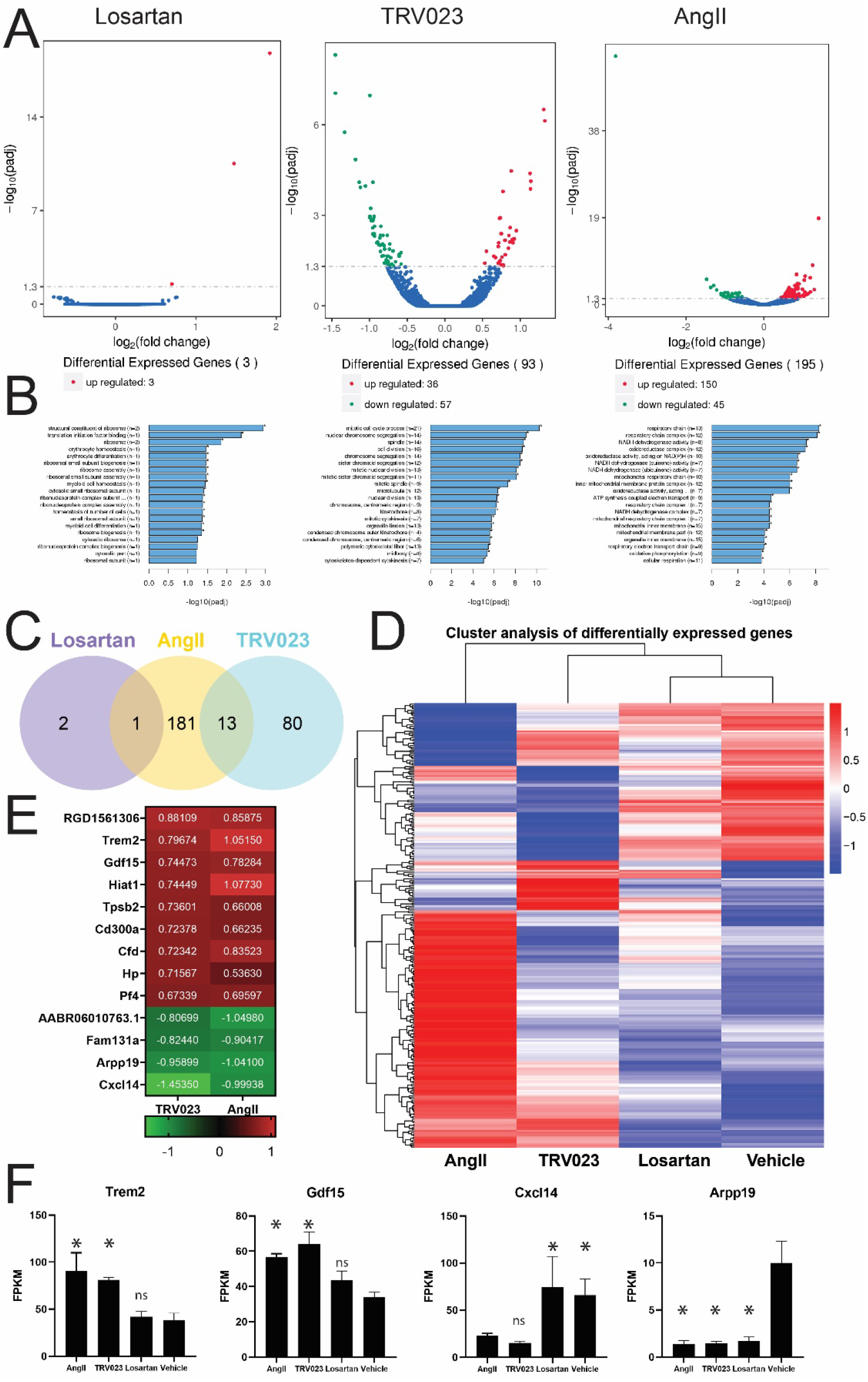
RNA sequencing of lungs from AT_1_R ligand-treated MCT rats demonstrates that TRV023 and AngII regulate pathways important in proliferation and metabolism. Total RNA from lung tissues from MCT rats treated with or without AngII, TRV023, losartan and vehicle were isolated and RNA sequencing was performed (n=3 rats per group). **A**, Volcano plot of differentially expressed genes between losartan, TRV023 and AngII from vehicle. **B**, GO pathways from those differentially expressed genes for losartan, TRV023 and AngII, respectively. **C**, Venn diagram of genes that were differentially expressed relative to vehicle in the losartan, TRV023 and AngII groups. **D**, Hierarchical clustering of differentially expressed genes between AngII, TRV023, losartan and vehicle. **E**, Heat map showing selected differentially expressed genes shared between AngII- or TRV023-treated MCT rats. **F**, Increased mRNA expression of MAPK signal regulating candidates *Trem2* and *Gdf15* by RNA sequencing. Decreased mRNA expression of negatively MAPK regulating candidates *Cxcl14* and *Arpp19* by RNA sequencing. Pathway enrichment performed by DAVID and GO enrichment tool. Statistical analysis was performed by one-way ANOVA. *p < 0.05 versus vehicle treated MCT rats. FPKM – fragments per kilobase of exon model per million reads mapped.

### β-arrestin-mediated AT_1_R signaling promotes pathways important in vascular proliferation and remodeling

Based on our transcriptomic findings, we hypothesized that the MAP kinase signaling cascade is activated in MCT PH rats treated with TRV023 or AngII. Previous studies have shown AngII-mediated transient activation of ERK at 2 to 5 minutes and remains elevated for 60 minutes in vascular smooth muscle cells (VSMCs) (34–36). To confirm if TRV023 also mediates activation of MAPK signaling in PASMCs from PH patients, we analyzed phosphorylation of ERK and p38 MAP kinases. PASMCs were grown at 60%-70% confluency and starved for 24 hours and treated with increasing concentration of AngII or TRV023 for 10 minutes for ERK phosphorylation and 20 minutes for p-38 phosphorylation. Immunoblots showed increased phosphorylation of ERK (**Fig. 4A** and **B**) and p38 (**Fig. 4A** and **C**) with increasing concentration of AngII as well as TRV023. The expression and activation of matrix metalloproteinases (MMPs) can also be of significant importance in vascular remodeling (8,37). To determine whether AT_1_R activation directly involves in the activation of matrix metalloproteinases (MMPs), consistent with our previous results, we found that AngII and TRV023 increased the mRNA expression of MMP-2 (**Fig. 4D**) but not tissue inhibitor of metallopeptidases 1 (TIMP-1) in PH PASMCs (**Fig. 4E**), consistent with an effect that would promote vascular remodeling. These data suggest β-arrestin-biased AT_1_R stimulation contributes to pulmonary vascular remodeling through multiple signaling axes.

**Figure 4.**
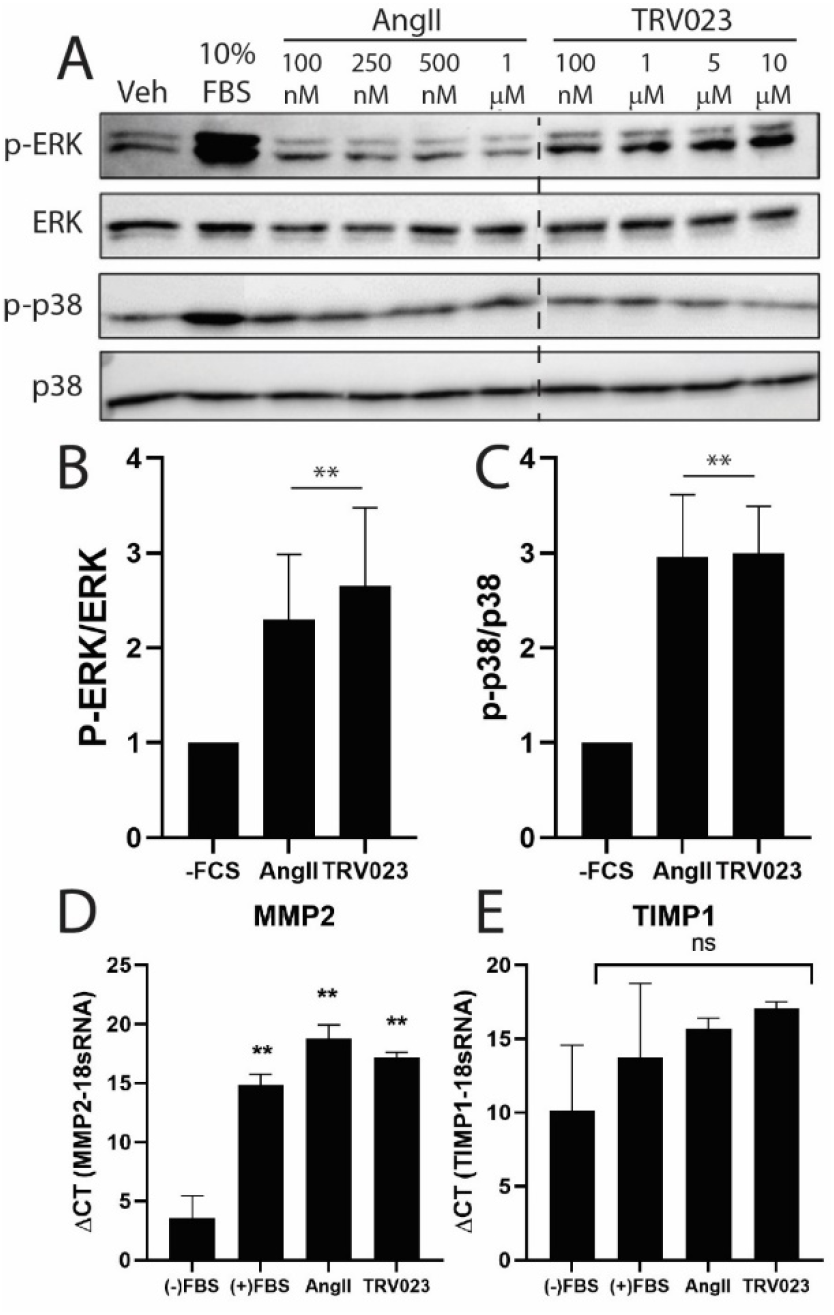
AngII and TRV023 activate MAP kinases in PASMCs isolated from PH patients. PASMCs isolated from PH patients were cultured and stimulated with AngII or TRV023 for analyzing phosphorylation of ERK and p38 at 10 and 20 mins respectively. (**A**) Representative immunoblot demonstrating AngII- and TRV023-induced phosphorylation of ERK and p38 activation in PASMCs isolated from PH patients. Quantitation of (**B**) phospho-ERK and (**C**) phospho-p38 MAP kinases from n=3 biological repeats of PH patients ([AngII] = 500 nM, [TRV023] = 5 μM). Real-time PCR of targets important in regulation of cell migration demonstrate that AngII and TRV023 both increase levels of (**D**) MMP2 with no significant expression of (**E**) TIMP1 expression. **, p < 0.05 by one-way ANOVA from vehicle treated samples.

### β-arrestin-mediated AT_1_R signaling stimulates human PASMC proliferation and migration

AngII stimulates VSMC proliferation and migration by activating tyrosine kinases, MAPK signaling and increasing intracellular Ca^2+^ levels that are regulated by both G proteins and β-arrestins downstream of the AT_1_R (38–40). To examine the effects of TRV023 on pulmonary artery SMC (PASMC) proliferation and migration *in vitro*, we performed bromodeoxyuridine (BrdU) proliferation and migration (wound healing scratch assays) in PASMCs isolated from PH patients. PH PASMCs were stimulated with AngII and TRV023 and proliferation monitored after stimulation for AngII (**Fig. 5A**) and TRV023 (**Fig. 5B**) (39). To examine the effects of AngII and TRV023 on PASMC migration, PASMCs were subjected to *in vitro* scratch wound healing assay (**Fig. 5C**) in the presence or absence of AngII, TRV023 or losartan. The scratch assay showed that both AngII and TRV023 increased PASMC migration, while losartan significantly decreased the rate of migration induced by both ligands (**Fig. 5D** and **E**). These findings demonstrate that AngII and TRV023 both promote PASMC proliferation and migration via the AT_1_R, consistent with a phenotype of pulmonary vascular remodeling *in vivo*.

**Figure 5.**
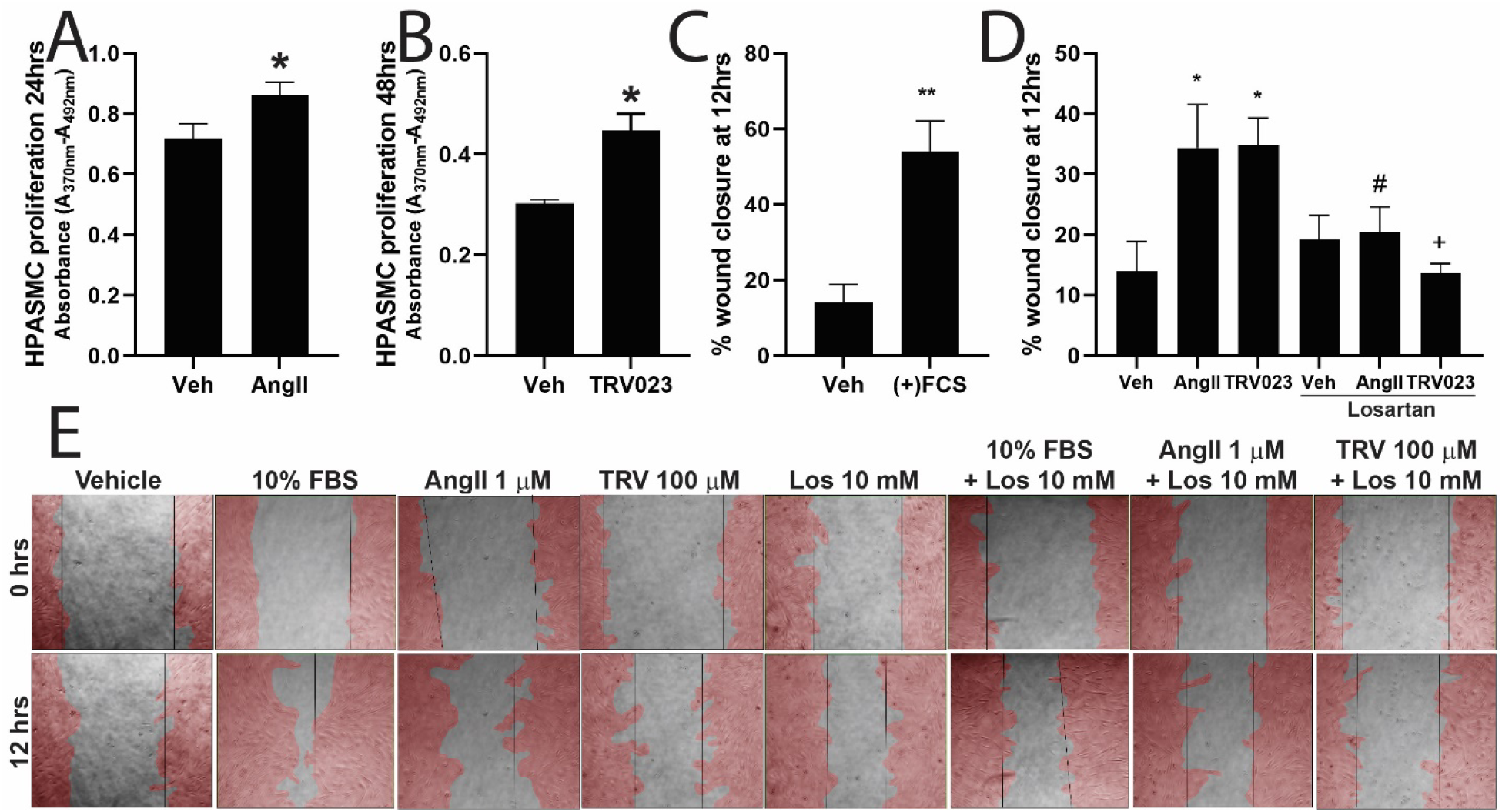
AngII and TRV023 promote PH-PASMC proliferation and migration. (**A**) AngII-mediated proliferation of PASMCs at 24 hours. (**B**) TRV023-mediated proliferation of PASMCs at 48 hours. Quantification of (**C**) FBS-induced and (**D**) AngII- and TRV023-induced PASMCs migration at 12 hours. AngII- and TRV023-mediated migration was blocked by losartan. (**D**) Representative live cell imaging of AngII and TRV023 induced PASMCs migration at 0 and 12 hours. Manual cell outlines are shown in red. Statistical analysis was performed by one-way ANOVA. *p < 0.05 versus vehicle, ^#^p < 0.05, ^+^p < 0.05 losartan treated versus non-treated. n=3 biological repeats of PH patients.

## Discussion

Despite current treatments, PAH is a devastating disease with a prognosis worse than many cancers and a 3-year survival of 68–70% (41,42). Therefore, new strategies are urgently needed to directly address the pathological pulmonary vascular remodeling that underlies PAH. Current PAH therapies such as prostacyclin receptor agonists and endothelin receptor antagonists act as pulmonary vasodilators through their effects on VSMCs (43,44). These agents have been proven to be effective in PAH, with benefits in exercise capacity and clinical endpoints (45–47). However, it is largely unknown to what extent the benefits of these drugs is related to their vasodilatory or antiproliferative properties. Here, we demonstrated that an AT_1_R β-arrestin-biased agonist that acts as a vasodilator (by blocking G protein-mediated signaling) while promoting signaling through β-arrestins did not have a beneficial effect and led to adverse pulmonary vascular remodeling and worse outcomes compared to losartan, an AT_1_R antagonist that blocked both G protein- and β-arrestin-mediated signaling (**Figure 6**). Unfortunately, we could not compare the responses of these drugs to strongly G protein-biased AT_1_R agonists, which have not been developed at this time (21). Similarly, the lack of biased agonists targeting the endothelin and prostacyclin receptors currently precludes a similar study of those receptors. Our findings suggest that the beneficial aspects of current PAH-specific therapies are not through acute effects on vasoconstriction but through their long-term effects on pulmonary vascular remodeling.

**Figure 6.**
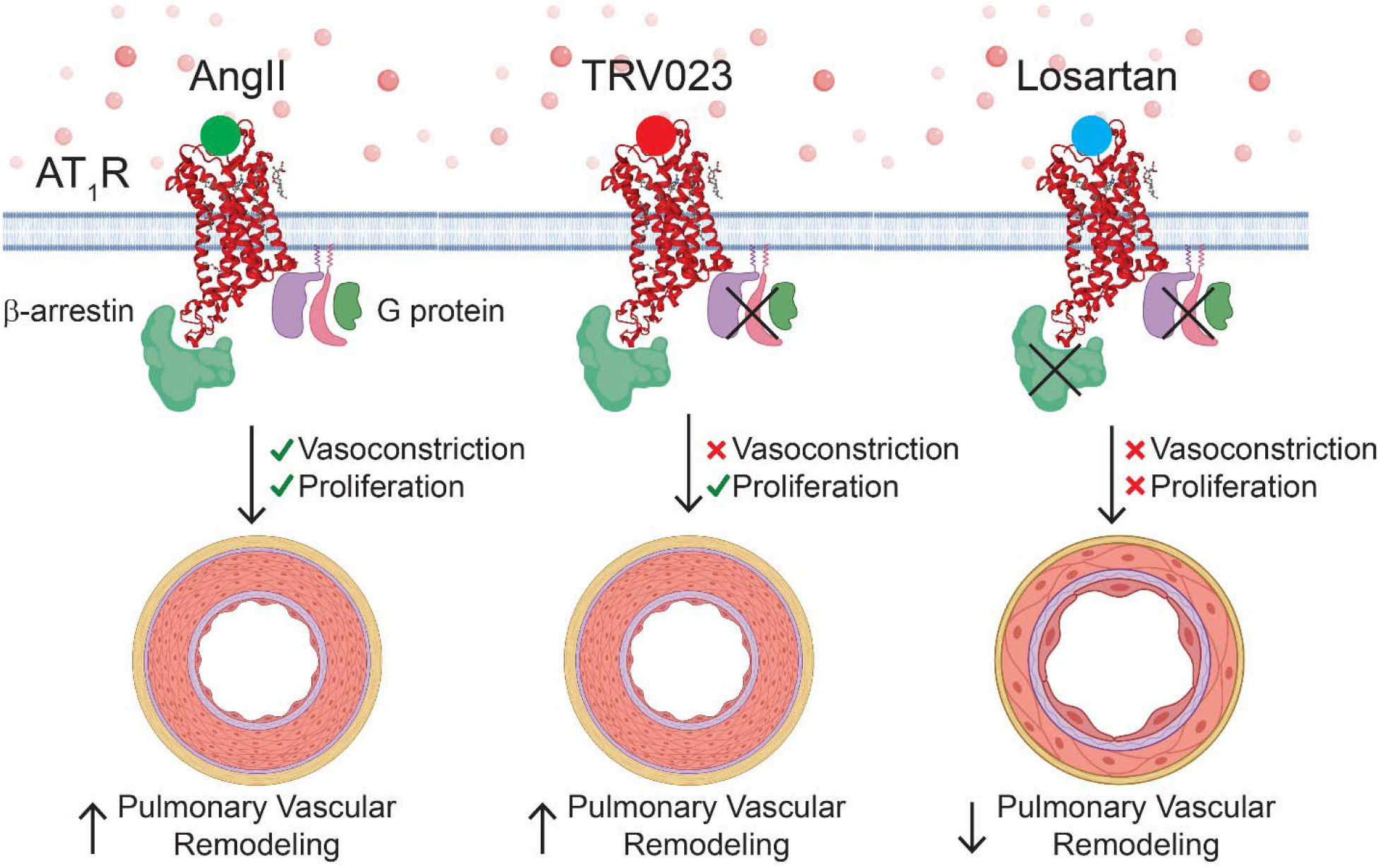
AT_1_R G protein- and β-arrestin-mediated signaling contribute to pulmonary vascular remodeling in PAH. With AngII, either through high circulating levels associated with the development of PH or through exogenous administration, there is G protein-mediated vasoconstriction as well as abnormal cellular proliferation, likely mediated by both G proteins and β-arrestins. TRV023 blocked G protein-mediated vasoconstriction but still promoted cellular proliferation and pulmonary vascular remodeling. Losartan blocked both G protein- and β-arrestin-mediated signaling, thereby decreasing both vasoconstriction and cell proliferation, resulting in less pulmonary vascular remodeling. Created with Biorender.com.

In contrast to our study in PAH, previous work has shown that β-arrestin-biased AT_1_R agonists have a number of distinct physiological effects compared to AngII in other models of cardiovascular disease. For example, acute infusion of AngII markedly increased mean arterial pressure in male spontaneously hypertensive rats, accompanied by a reduction of glomerular filtration rate (46), while TRV023 blocked the acute infusion of AngII-induced hypertension state in a dose-dependent manner in mice (16). Likewise, chronic administration of AngII in rats has been shown to induce LV hypertrophy (24), which can be blunted by co-administration of TRV023 or losartan (24). Consistent with these results, we found that acute infusion of AngII, but not of TRV023, increased LV and RV systolic pressures. The β-arrestin-biased agonist TRV120027 (15) competitively antagonized AngII-induced mean arterial pressure elevation while stimulating cardiac contractility as determined by load-independent measurements (15,24), whereas the AT_1_R antagonist telmisartan decreased mean arterial pressure and cardiac contractility (15). TRV120027 has also been shown cause a neonatal-specific sustained positive inotropic effect without increasing heart rate, an effect thought to be mediated by its activation of L-type calcium channels (48). TRV023 has been shown to increase LV contractility and promote cell survival in mice, the effects of which are dependent on β-arrestin (16). Taken together, conventional and β-arrestin-biased AT_1_R agonists are pharmacologically distinct *in vivo*.

In preclinical models of left heart failure, TRV120027 decreased mean arterial pressure, right atrial pressure, pulmonary capillary wedge pressure, and systemic and renal vascular resistances, while increasing cardiac output and renal blood flow (18,49). In a mouse model of familial dilated cardiomyopathy, TRV023 increased cardiac performance, suggesting that AT_1_R biased ligands may prove to be a novel inotropic approach in familial dilated cardiomyopathy (17). Furthermore, in line with the preclinical findings, infusion of TRV120027 was safe and well tolerable and leads to the reductions of mean arterial pressure in healthy volunteers with sodium intake restriction (50). In contrast to left heart failure, PAH is characterized by a marked increase in RV afterload and subsequent dilatation, fibrosis and RV failure. The mechanisms underlying the development of RV failure secondary to PAH remain an area of active investigation. RV function determines the symptoms of patients with PAH and is one of the most important factors for survival. Unlike LV dysfunction, there are currently no therapies that directly target the RV in PAH (51,52). Here, we found that losartan significantly improved survival compared to vehicle, AngII or TRV023. In contrast to its effects in models of left ventricular dysfunction, TRV023 failed to prolong the survival compared to control or AngII, and TRV023 did not improve the hemodynamics or enhance cardiac performances in RV PV loop analyses. These results suggest that β-arrestin-biased AT_1_R ligands are not beneficial for targeting the RV or pulmonary vasculature in PAH. Our observations are consistent with a previous study that showed no differences in RV hypertrophy between AT_1_R antagonist losartan and vehicle-treated rats with MCT-induced PAH (4). However, in contrast to our findings, they observed RV afterload was reduced in losartan-treated rats with PAH, without affecting RV contractility (4). This inconsistency may be due to differences in treatment and follow-up time.

The lack of cardiac effects of TRV023 led us to examine potential AT_1_R-mediated pulmonary vascular remodeling. Pulmonary vascular remodeling in PAH is characterized by endothelial dysfunction (27), clonal expansion of apoptosis-resistant endothelial cells, and smooth muscle cell proliferation, along with a complex interplay between adventitial fibroblasts, perivascular inflammatory cells, and the extracellular matrix (1,53). In MCT-induced PAH rats, there were no discernible changes in PA wall media thickness across different treatments. Intriguingly, TRV023 stimulated cellular proliferation in pulmonary arteries as determined by Ki67 compared to vehicle, but not AngII. This disparity could be explained by increased apoptosis as quantified by the elevation of cleaved caspase-3 in AngII-treated pulmonary arteries, consistent with a previous study that showed that AngII markedly increased proapoptotic activity in the suprarenal aortas in *ApoE* knockout mice (54). Our transcriptomic data demonstrated that the TRV023 group had higher levels of transcripts associated with cell proliferation. These results are consistent with TRV023 promoting pulmonary vascular remodeling in MCT-induced PAH rats.

AT_1_R overexpression in patients with idiopathic PAH correlates with the activation of MAPK and SRC signaling compared to controls (4) and AngII treatment selectively induces the proliferation of smooth muscle cells isolated from patients with idiopathic PAH (4). Consistent with this, AngII and TRV023 activated ERK1/2 and SRC signaling in U2OS and HEK cells overexpressing rat AT_1_R (15). Here we found that a number of signaling effectors regulated by MAPK signaling pathways were increased in MCT rats treated with AngII or TRV023 compared to vehicle or losartan in the lung transcriptome analysis. We further verified our results *in vitro* that AngII and TRV023 stimulated the proliferation and migration of PASMCs and induced the phosphorylation of ERK1/2 and p38 in PH PASMCs, effects that were blunted by losartan. As SMC proliferation and migration can be induced by alterations in the composition of the extracellular matrix from MMP activation (37), we then tested the effects of AngII and TRV023 on transcript levels of MMP-2 and TIMP-1 in PH PASMCs. We found that both AngII and TRV023 increased MMP-2 levels, likely through the activation of Erk1/2 signaling by AngII and TRV023, as it has been demonstrated that tryptase activates the Erk1/2 signaling cascade to induce the production of fibronectin and MMP-2 in PASMC (55). Taken together, β-arrestin-biased ligands likely promote PASMC proliferation and migration through multiple pathways, contributing to pulmonary vascular remodeling in pulmonary hypertension.

In conclusion, our findings demonstrate that activating β-arrestin-biased AT_1_R signaling promotes vascular remodeling and is undesirable in PAH. This suggests that the development of new PAH therapies at targets that have previously only been considered to only promote vasodilation, such as GPCRs, can still reverse pulmonary vascular remodeling in PAH through mitigating apoptosis resistance and angioproliferative lung pathology.

## Author Contributions

ZM, GV and SR designed the study and wrote the manuscript. ZM performed and analyzed the experiments for chronic infusion, survival studies and IF. GV, XY and LM performed the animal experiments for acute infusion. GV analyzed data for Ki67 and cleaved caspase-3 staining. GV, MS, NN and IC performed *in vitro* experiments. SR analyzed data for RV PV loop and supervised the experiments. All authors reviewed the results and approved the final version of the manuscript.

## Acknowledgements

We thank Lan Mao in the Duke Cardiovascular Research Center Physiology Core for help with right and left heart catheterization, Zuowei Su, Ph.D. at Research Immunohistology Lab in the Department of Pathology, Duke University Medical Center for Ki67 and cleaved caspase-3 staining, and Yasheng Gao in the Duke Light Microscopy Core Facility for help with live cell imaging. SR was funded by a Gilead Research Scholars in Pulmonary Hypertension, Burroughs Wellcome Career Award for Medical Scientists Award, and K08HL114643.

