## Supplemental data for "β-arrestin-mediated Angiotensin II type 1 Receptor Activation Promotes Pulmonary Vascular Remodeling in Pulmonary Hypertension"

**Running Title:** β-arrestin-mediated $AT_1R$ signaling promotes PAH


Zhiyuan Ma[1†], Gayathri Viswanathan[1†], Mason Sellig[2], Chanpreet Jassal[4], Issac Choi[1], Xinyu Xiong[1], Nour Nazo[1], Sudarshan Rajagopal[1, 3, *]

[1]Division of Cardiology, Department of Medicine, Duke University School of Medicine, Durham, NC, USA

[2]Trinity College of Arts and Sciences, Duke University, Durham, NC, USA.

[3]Department of Biochemistry, Duke University Medical Center, Durham, NC, USA.

[4]The University of North Carolina, Chapel Hill, NC, USA.

[†]Z. Ma and G. Viswanathan contributed equally to this work

*To whom correspondence should be addressed:

Sudarshan Rajagopal, MD, PhD

Division of Cardiology, Department of Medicine, Duke University Medical Center

160B CARL Building, 130 Research Drive, Durham, NC  27710

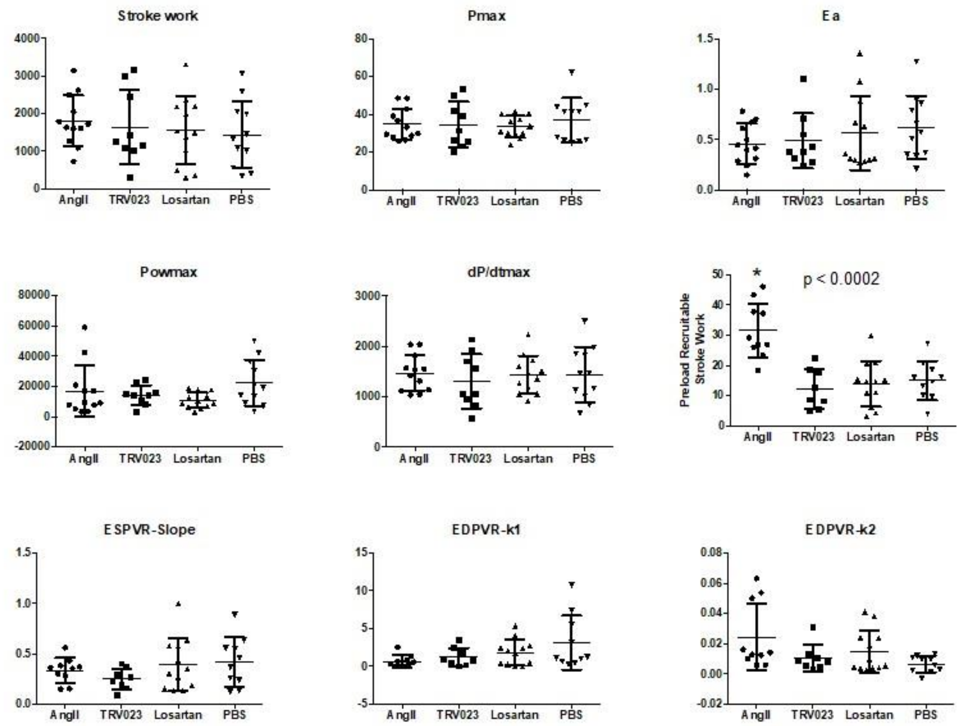


**Supplemental Figure 1. Hemodynamic effects of chronic infusion of $AT_1R$ ligands on right ventricular pressure-volume loop parameters in MCT PH rats.** Chronic infusion of AngII, TRV023 or losartan in MCT rats resulted in no significant changes in the majority of hemodynamic parameters from pressure-volume loop analysis of the right ventricle. The only significant difference that was noted was an increase in preload recruitable stroke work in AngII-treated rats ($p < 0.\ 0002$). Maximal pressure (Pmax), End-systolic elastance (Ea), Maximal power (Pmax), Maximum of Change in Pressure / Change in time (dP/dtmax), end-systolic pressure volume relationship slope (ESPVR-slope), end-diastolic pressure volume relationship coefficient 1 (EDPVR-k1), end-diastolic pressure volume relationship coefficient 2 (EDPVR-k2). Statistical analysis was performed by one-way ANOVA and Turkey's multiple comparisons test.

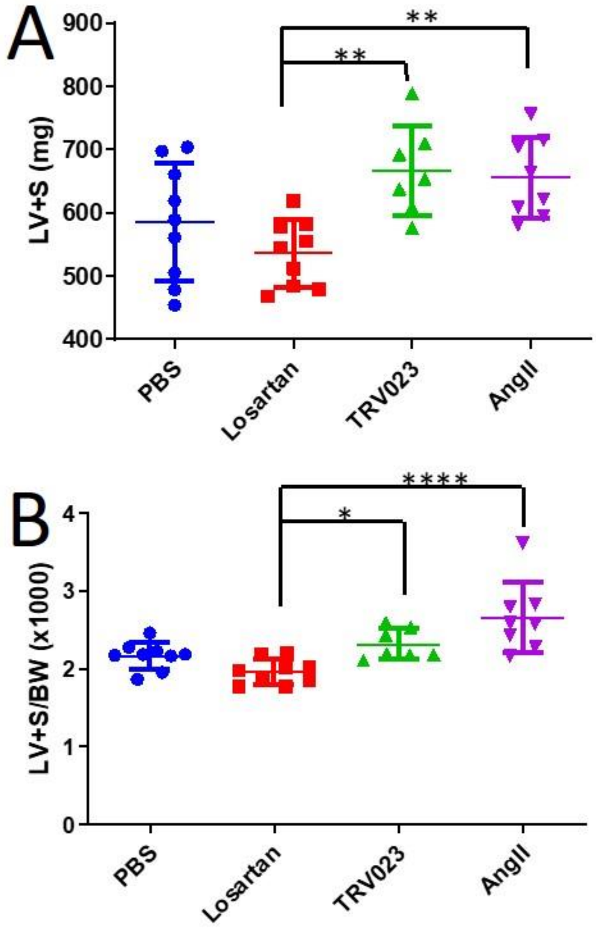


**Supplemental Figure 2. Effects of chronic infusion of AT1R ligands on LV hypertrophy in MCT rats.** Treatment with TRV023 and AngII both induced LV hypertrophy compared to losartan. (**A**) as assessed by left ventricular + septum (LV+S) weight and (**B**) LV+S corrected by rat body weight. Statistical analysis was performed by one-way ANOVA and Turkey's multiple comparisons test. (*, $p < 0.05$, **, $p < 0.01$, ****, $p < 0.0001$)

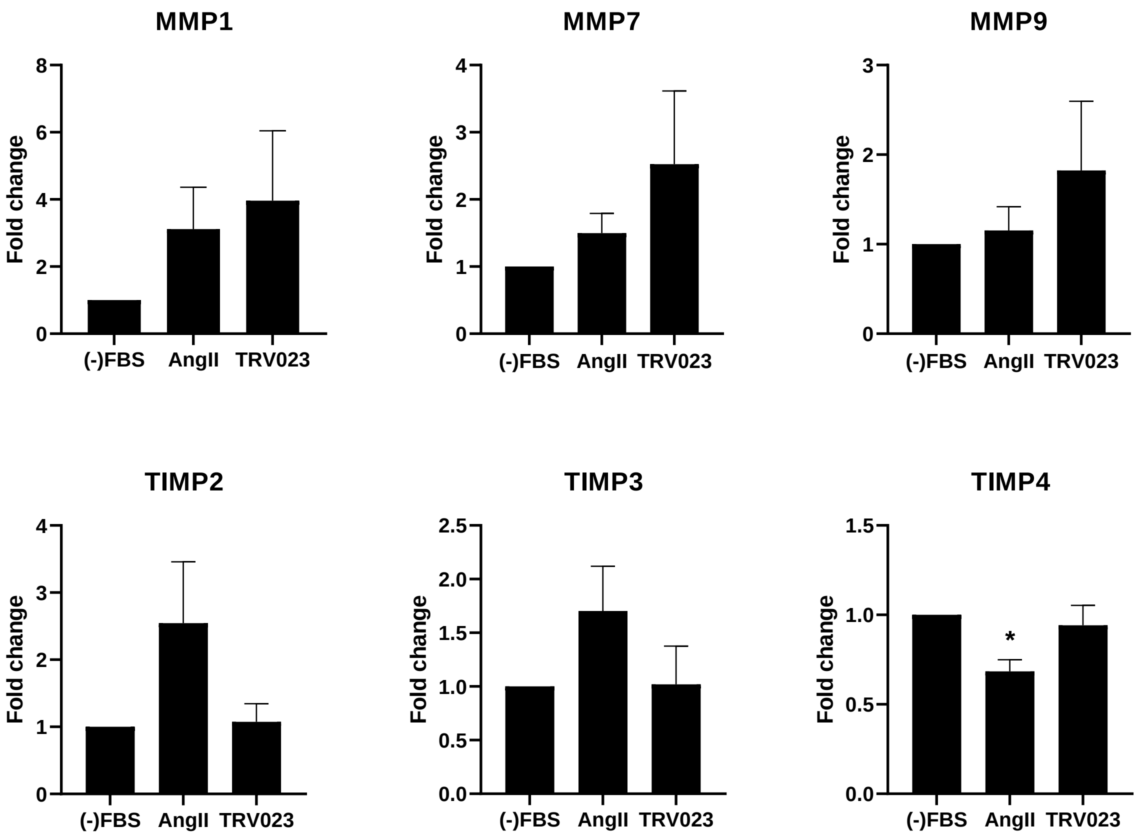


**Supplemental Figure 3. AngII and TRV023 alter the composition of the extracellular matrix in PASMCs isolated from PH patients.** PASMCs isolated from n=3 biological repeats of PH patients were cultured and stimulated with 500nM AngII or 5 μM TRV023 for analyzing mRNA expression of MMPs and TIMPs. *, $p < 0.05$ by one-way ANOVA from vehicle treated samples.

**Table 1. List of PCR primers**

| Gene | Forward primer | Reverse primer |
|---|---|---|
| MMP-9 | GCACGA CGT CTT CCA GTA CC | CAG GAT GTC ATA GGT CAC GTA GC |
| MMP-7 | GTATGGGACATTCCTCTGATCC | CCAATGAATGAATGAATG GATG |
| MMP-1 | CCTAGCTACACCTTCAGTGG | GCCCAGTACTTATTCCCTTT |
| MMP-2 | ATGACAGCTGCACCACTGAG | CTCCTGAATGCCCTTGATGT |
| MMP-9 | TACCCTATGTACCGCTTCAC | GAACAAATACAGCTGGTTCC |
| 18S rRNA | GTAACCCGTTGAACCCCATT | CCATCCAATCGGTAGTAGCG |
| TIMP-1 | TGACATCCGGTTCGTCTACA | GTTTGCAGGGGATGGATAAA |
| TIMP-2 | CCGCAACAGGCGTTTTGCAA | TCACTTCTCTTGATGCAGGC |
| TIMP-3 | TTCTGCAACTCCGACATCGT | ATGCAGGCGTAGTGTTTGGA |
| TIMP-4 | CACTACCATCTGAACTGTGGCTG | GCTTTCGTTCCAACAGCCAGTC |

**Table 2. List of abbreviations**

| | |
|---|---|
| PAH | Pulmonary arterial hypertension |
| RV | Right ventricular |
| AT1R | Angiotensin II type 1 receptor |
| GPCR | G protein-coupled receptor |
| TRV023 | TRV120023 |
| MCT | Monocrotaline |
| LV | Left ventricular |
| PASMCs | Pulmonary artery smooth muscle cells |
| ETAR | Type A endothelin receptor |
| IP | Prostacyclin receptor |
| $Ca^{2+}$ | Calcium |
| PKC | Protein kinase C |
| MAPK | Mitogen-activated protein kinase |
| ERK | Extracellular signal-regulated kinases |
| AngII | Angiotensin-2 |
| SD | Sprague–Dawley |
| EDRVP | End-Diastolic Right Ventricular Pressure |
| EDLVP | End-Diastolic Left Ventricular Pressure |
| RVPmax | Peak systolic RV pressure |
| LVPmax | Maximal left ventricular systolic pressure |
| IVC | Inferior vena cava |
| PV | Pressure-volume |

| | |
|---|---|
| RV/BW | Right ventricular to body weight |
| (LV+S)/BW | Left ventricular and septal weight to body weight |
| VWF | Von Willebrand factor |
| Ki67 | Cellular marker for proliferation |
| HBSS | Hank's balanced salt solution |
| HEPES | 4-(2-hydroxyethyl)-1-piperazineethanesulfonic acid |
| SMC | Smooth muscle cell |
| ELISA | Enzyme-linked immunosorbent assay |
| BrdU | Bromodeoxyuridine |
| PBS | Phosphate-buffered saline |
| SDS-PAGE | Sodium dodecyl sulfate-polyacrylamide gel electrophoresis |
| BSA | Bovine Serum Albumin |
| RNA | Ribonucleic acid |
| DEG | Differential gene expression analysis |
| DAVID | Database for Annotation, Visualization and Integrated Discovery |
| qPCR | Quantitative polymerase chain reaction |
| cDNA | Complementary Deoxyribonucleic acid |
| MMP | Matrix metalloproteinases |
| rRNA | Ribosomal ribonucleic acid |
| TIMP | Tissue inhibitor of metallopeptidases |
| ANOVA | Analysis of variance |

| | |
|---|---|
| dP/dt | Change in pressure over change in time |
| RVSP | Right ventricular systolic pressure |
| RNA-seq | RNA sequencing |
| GO | Gene ontology |
| PANTHER | Protein analysis through evolutionary relationships |
| Trem2 | Triggering receptor expressed on myeloid cells 2 |
| Gdf15 | Growth differentiation factor 15 |
| p-38 | Mitogen activated protein kinase 14 |
| JNK | Mitogen activated protein kinase 8 |
| Arpp19 | cAMP-regulated phosphoprotein 19 |
| VSMC | Vascular smooth muscle cell |
| PA | Pulmonary arteries |
| ApoE | Apolipoprotein E |
| SRC | Proto-oncogene tyrosine-protein kinase |
| U2OS | Human osteosarcoma cell line |
| HEK | Human embryonic kidney 293 cells |
| S | Septum |
| SEM | Standard error mean |
| Pmax | Maximal pressure |
| BW | Body weight |
| LV+S | Left ventricular + septum |
| EDPVR-k2 | End-diastolic pressure volume relationship coefficient 2 |

| | |
|---|---|
| EDPVR-k1 | End-diastolic pressure volume relationship coefficient 1 |
| ESPVR-slope | End-systolic pressure volume relationship slope |
| dP/dtmax | Maximum of Change in Pressure / Change in time |
| Pmax | Maximal power |
| Ea | End-systolic elastance |
